# Voxelwise encoding models with non-spherical multivariate normal priors

**DOI:** 10.1101/386318

**Authors:** Anwar O. Nunez-Elizalde, Alexander G. Huth, Jack L. Gallant

## Abstract

Predictive models for neural or fMRI data are often fit using regression methods that employ priors on the model parameters. One widely used method is ridge regression, which employs a spherical Gaussian prior that assumes equal and independent variance for all parameters. However, a spherical prior is not always optimal or appropriate. There are many cases where expert knowledge or hypotheses about the structure of the model parameters could be used to construct a better prior. In these cases, non-spherical Gaussian priors can be employed using a generalized form of ridge known as Tikhonov regression. Yet Tikhonov regression is only rarely used in neuroscience. In this paper we discuss the theoretical basis for Tikhonov regression, demonstrate a computationally efficient method for its application, and show several examples of how Tikhonov regression can improve predictive models for fMRI data. We also show that many earlier studies have implicitly used Tikhonov regression by linearly transforming the regressors before performing ridge regression.

## 1 Introduction

Cognitive and systems neuroscience has in recent years become increasingly reliant on predictive encoding models. In the fMRI literature, encoding models have produced insights into the cortical representations of visual (Thirion et al., 2006, Kay et al., 2008b, Nishimoto et al., 2011, Huth et al., 2012), auditory (De Angelis et al., 2017, de Heer et al., 2017), and linguistic (Mitchell et al., 2008, Wehbe et al., 2014, Huth et al., 2016) information. To efficiently estimate the parameters of encoding models, many studies use L2-regularized (ridge) regression (Hoerl and Kennard, 1970). L2 regularization improves regression models by imposing a multivariate normal prior on the model parameters, where the mean of the prior is zero and the covariance is spherical. Compared to unregularized regression, ridge makes models better at generalizing to new data and decreases overfitting by shrinking model parameter estimates towards zero and improving estimation for features that are nearly collinear. However, assuming a spherical covariance is rarely optimal, and in many cases prior information or expert knowledge can be used to construct informative non-spherical priors. In this paper we explore how non-spherical priors can be applied to several encoding model problems and show that this can greatly improve model performance. We also show that some previously published encoding models can be reinterpreted in terms of non-spherical priors, providing new insights into why those models were successful. Finally, we offer practical advice and efficient methods for estimating encoding models with non-spherical priors.

Although encoding models have proven highly successful for modeling fMRI data, there are several complications that make them difficult to use. First, in many feature spaces it is difficult to assign a specific interpretation to the features. This problem is especially acute for feature spaces learned using unsupervised methods, such as the word embedding space word2vec (Mikolov et al., 2013). When using these feature spaces to predict neural or BOLD responses, it is difficult to interpret what exactly a given voxel represents.

Second, it is often unclear how the regularization method used for regression interacts with the choice of feature space. For example, feature spaces that are identical up to a linear transformation (i.e. 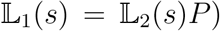) can yield drastically different results even though both span the same space.

Third, although the basic shape and variability of the HRF are reasonably well understood (Boynton et al., 1996, Büchel et al., 1998, Friston et al., 1998, Glover, 1999), many studies do not use this prior information when estimating the HRF (Kay et al., 2008a, Nishimoto et al., 2011, Huth et al., 2012), and most studies simply assume a single canonical HRF for all voxels (Penny et al., 2011).

Fourth, it is becoming increasingly important to characterize how and where different feature spaces overlap in terms of variance explained (Lescroart et al., 2015, de Heer et al., 2017). This is usually done by combining different feature spaces into one encoding model. However, ordinary regularization techniques wrongly assume that all feature spaces require the same level of regularization.

Here we address all of these issues by constructing encoding models using carefully designed multivariate normal priors. In the standard encoding model formulation, complex features are extracted from the stimuli and then regularized regression is used to learn model parameters subject to simple priors. In the new framework presented here, we extract simple, interpretable features from the stimuli, and then use Tikhonov regression (Tikhonov et al., 1977) to learn parameters subject to complex multivariate priors. This is made possible by a duality between imposing a prior and extracting features from the stimuli, so the exact same model can be represented in both ways. This simple change in perspective has significant consequences for model interpretation, because it shows that a complex feature space can be decomposed into a combination of a simple feature space and a multivariate prior. This framework is also highly modular, making it easy to combine different spatial and temporal priors and test many different kinds of priors.

We evaluate each proposed application of our framework on empirical data from naturalistic experiments on vision and language. We show that non-spherical multivariate normal priors can improve prediction accuracy in a variety of settings. In order to encourage the adoption of the framework presented here, we have released an open-source Python software package (http://github.com/gallantlab/tikreg) that efficiently implements all the models described in this paper.

## 2 Linearized predictive encoding models

In a typical fMRI experiment, brain images 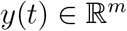 are recorded at times *t* =1& *T* while a subject is exposed to stimuli *s*(*t*) (Figure 1). Each brain image consists of m voxels, *y*ℓ(*t*) for ℓ = 1… *m*. The goal of the encoding model framework is to find a function that maps stimuli to BOLD responses in each voxel: *f*ℓ(*s*(1),…, *s*(*t*)) ≈ *y*ℓ(*t*). Because the space of possible functions is extremely large, it is common to work under a hypothesis that limits the complexity of *f*. Although there often are many reasonable hypotheses that one can make about *f*, the only type that we shall consider here is where *f* is a linear combination of features that are extracted from the stimulus, usually by a nonlinear function. In this case, *f* is called a “linearized” model, and the function that extracts features from the stimulus, 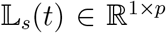, is called the “linearizing transformation” (Wu et al., 2006). Formally, 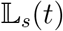 maps a *u*-dimensional stimulus at time *t* into a *p*-dimensional vector of stimulus features *X_i_*(*t*).

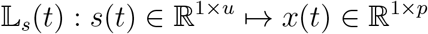

**Figure 1.**
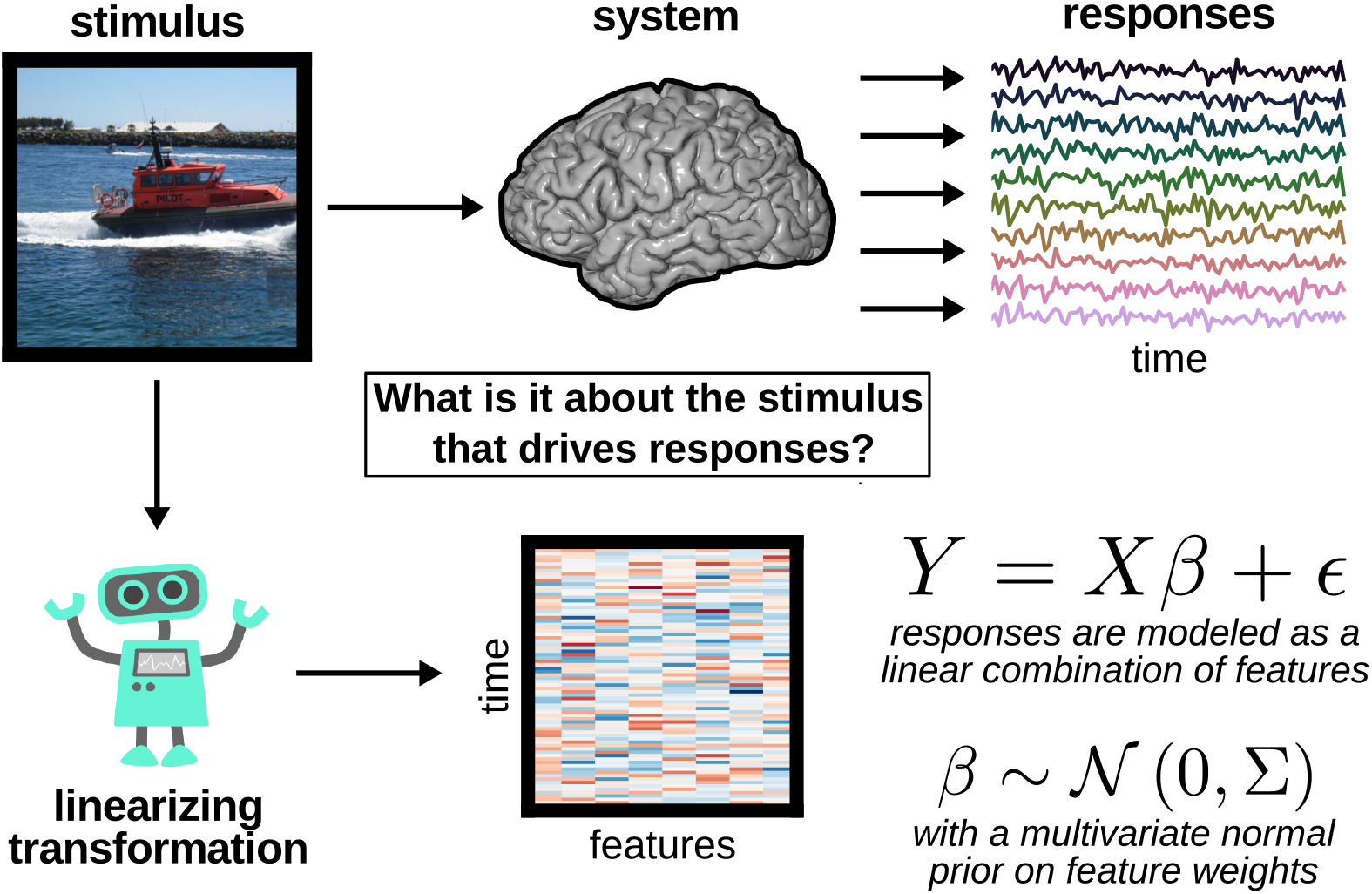
Modeling the stimulus-response relationship with linearized encoding models and multivariate normal priors. In a typical fMRI experiment a series of stimuli are shown to a subject while their brain activity is recorded. In the voxelwise encoding model framework, features are first extracted from the stimuli using a computational model, human labels, or by any other method. Brain activity is then modeled as a weighted, linear combination of the features. Some form of regularization is usually required when a large number of features are used or when the signal-to-noise ratio is low. The most common method of regularization is to impose a prior distribution on the feature weights. When the distribution is a spherical multivariate normal (MVN), this is called ridge regression. A spherical MVN distribution assumes equal variance in all dimensions and zero covariance. More generally the MVN prior can also have non-spherical structure. This is called Tikhonov regression.

Under the linearized model formulation, the brain response is modeled as a linear combination of the stimulus features, usually over a fixed time window *d*,

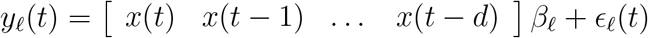

where 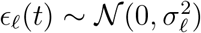 is stationary, zero-mean normal noise, 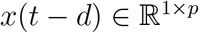 is the feature vector delayed *d* time points, and 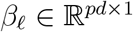 is a set of linear weights over the *p* features at each of the *d* delays.

To write the simultaneous equation for all voxels we replace *yℓ*(*t*) with a matrix 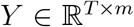 that contains the response of each voxel at each timepoint, and we replace βℓ with a matrix 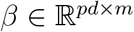 that contains the weight vector for every voxel. We write the matrix of linearized stimulus features as 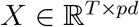,

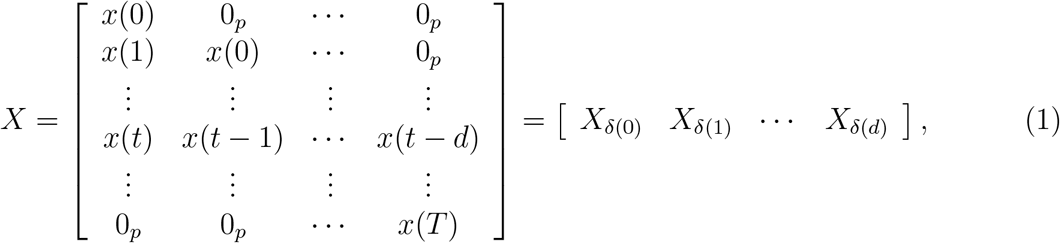

where each row of *X* contains the feature vectors for the past *d* timepoints. Each block of *p* columns *X_δ_*(*j*) contains the linearized stimulus feature matrix delayed by *j* time points. This is referred to as a finite impulse response model (Oppenheim et al., 1983). This allows us to rewrite the basic model as:

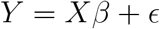

where 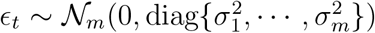 is zero-mean, independent noise for each voxel at each time point. The only free parameter in this formula is the weight matrix 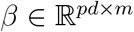.

We can find an estimate of *β* by maximizing the probability of the data *Y* given the stimulus features *X*

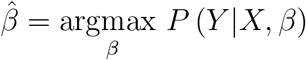

This estimate of β is called the maximum a posteriori (MAP) estimate. We can derive various analytic solutions depending on the form of the distribution we assume for *P*(*Y|X, β*). In this paper, we assume that the responses can be modeled as multivariate normal random variables.

The likelihood of the data can be expressed as

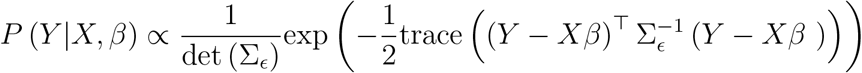

where 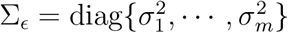 contains the variance of the noise for each voxel. If we assume that the noise variance is the same in each voxel, then we can set Σ_∊_ = σ^2^*I*. We can also switch to using the log of the likelihood instead of the likelihood. The log-likelihood of the data can then be expressed as

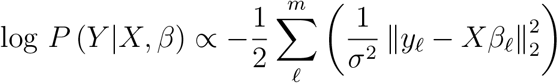

Finally, note that maximizing the log-likelihood is equivalent to minimizing the negative log-likelihood of the data. The estimate for all voxels can be found simultaneously by solving

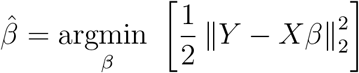

which is equivalent to finding the *β* that minimizes the squared difference between predicted and actual responses (i.e. ordinary least squares regression).

However, finding the value of *β* that exactly minimizes this squared error function often produces results that do not generalize to new stimuli. This is due to overfitting. Overfitting occurs when model parameters capture the noise e in addition to the underlying signal. This is a common problem when the data available to estimate the model parameters is small. When building a predictive encoding model our goal is not simply to explain the data that is given, but to predict new data.

### 2.1 Ridge regression

To avoid overfitting it is common to employ regularized regression techniques (Friedman et al., 2001). Regularization imposes a prior distribution on β. This prior limits how well a model can explain the given data. The goal becomes to maximize the probability of the observed data, by finding the β that maximizes the product of the likelihood and the prior

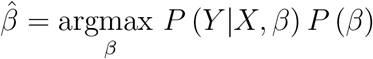

One commonly used regularization technique is ridge regression (Hoerl and Kennard, 1970), which imposes a zero-mean multivariate normal prior on the individual voxel weights *β*ℓ:

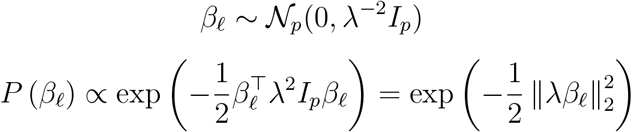

Ridge regression can be implemented by adding a penalty term to the error function, where the penalty is proportional to the sum of the squared weights,

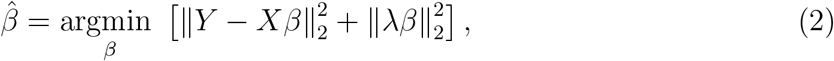

and the strength of the regularization is controlled by λ, the regularization coefficient. The closed-form solution for the ridge regression problem is given by

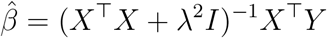

### 2.2 Tikhonov regression

Ridge regression imposes a zero-mean, spherical multivariate normal prior on the feature weights. However, expert knowledge can be used to create a more sophisticated, non-spherical multivariate normal prior on the weights,

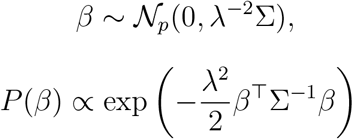

where 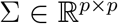 is the positive semidefinite prior covariance matrix. Note that λ is present as a scaling factor on Σ. This determines how much influence the prior has on the estimated weights.

If we factorize the inverse of the prior covariance matrix by taking its matrix square root Σ_−1_ = *C^⊤^ C*, where 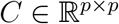 then

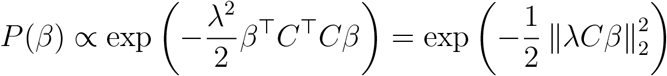

The problem can then be solved by maximizing the product of the likelihood and this new prior, or, as above, by minimizing the negative log likelihood,

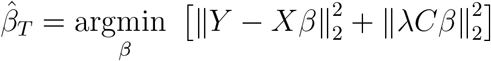

This is known as Tikhonov regression (Tikhonov et al., 1977). Here C can be thought of as a penalty matrix that punishes β when it does not conform to the prior. However, since there are many matrix square roots, C is not uniquely determined by the prior covariance, and in fact any C that satisfies the given relation will produce the same *β̂_T_*. Also note that when *C = I_p_* Tikhonov regression reduces to ridge regression (Hoerl and Kennard, 1970).

The Tikhonov minimization problem has a closed form solution,

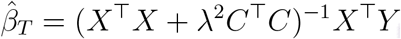

However, this formulation does not immediately admit efficient computational solutions, making it less useful for solving large-scale problems. Fortunately there is a computationally efficient method for solving Tikhonov regression problems. This method, which is often referred to as the “standard form” (Hansen, 1998), transforms a Tikhonov problem into a ridge regression problem. This transformation is accomplished in three steps.

First, a linear transformation is applied to *X*, giving

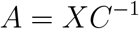

Second, ridge regression is carried out with A, giving

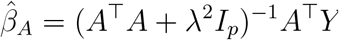

Third, the estimated weights are projected back into the original space to give the Tikhonov estimate,

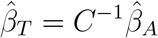

(For a proof of this see Appendix A). Because the standard form uses ridge regression internally, it is clear that Tikhonov regression in the standard form will admit the same efficient computational solutions as ridge regression.

The standard form transformation can be used to convert any Tikhonov regression problem into a ridge regression problem by way of a linear transformation of *X*. By the same logic, any linear transformation of *X* followed by ridge regression is equivalent to some Tikhonov regression problem, and thus some non-spherical multivariate prior on the model weights. This relationship has interesting implications for a number of neuroimaging studies that have applied ridge regression to linearly transformed stimuli, because the models employed by those studies can be re-interpreted as Tikhonov regression with non-spherical priors. We use this technique to explore and re-interpret the models used in some previous studies.

## 3 Using multivariate normal priors

### 3.1 Feature priors

### 3.1.1 Word embeddings

Several earlier studies have used word embedding spaces to model how the brain represents the meaning, or semantic content, of words (Mitchell et al., 2008, Wehbe et al., 2014, Huth et al., 2016). In this approach, each word is converted into a vector with anywhere from 20 (Mitchell et al., 2008) to 1000 (Huth et al., 2016) embedding dimensions. These vectors are constructed using word co-occurrence statistics from large corpora of text (Turney and Pantel, 2010), and are designed such that words with similar or related meanings (such as ‘month’ and ‘week’) are assigned similar vectors, but words with dissimilar meanings (such as ‘month’ and ‘tall’) are not. After converting words to vectors, regression models are used to predict BOLD responses as a function of the embedding dimensions.

Formally, this approach starts by defining a matrix of word indicator variables 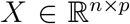, where *X_t_,_i_* = 1 if word *i* was presented at time *t* and 0 otherwise. Here n is the total number of time points and *p* is the total number of words in the experiment. Then, in order to replace each word with its *q*-dimensional embedding vector, the indicator matrix is multiplied with an embedding matrix 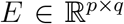 whose rows contain the word embedding vectors. Finally, regression is performed in the embedding space, yielding the linear model

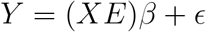

Interestingly, this formulation appears identical to the standard form transformation of Tikhonov regression (see Appendix A). If the model weights, *β*, are estimated using ridge regression (Wehbe et al., 2014, Huth et al., 2016), then this approach is equivalent to Tikhonov regression where the features are word indicators (i.e. the feature matrix is *X*), and the prior covariance is given by dot products between embedding vectors, Σ = *EE^⊤^* (and *C*^−1^ = *E*).

Thus, the word embedding approach is equivalent to putting a multivariate normal prior on the model weights across words, such that the prior covariance between weights for different words is equal to the dot product between their embedding vectors. If words that have similar meanings have similar embedding vectors, then the dot product between those vectors will be high, and the weights for those words will covary strongly. This re-interpretation of the word embedding approach seems in many ways to be more natural and intuitive than thinking of it as regression in the word embedding space, which is highly abstract and difficult to explain.

### 3.1.2 Evaluating feature MVN priors

To illustrate the Tikhonov approach to word embeddings we estimated two different linear models using the data from Huth et al., 2016. Both models use words as features, but one model applies an identity prior to the weights while the other applies a semantic similarity prior based on a word embedding space. The data come from an experiment where two subjects listened to approximately two hours of naturally spoken narrative stories while undergoing continuous BOLD fMRI. The stories were transcribed and then the transcripts were aligned to the audio to determine exactly when each word was spoken. These aligned transcripts were then used to generate the word indicator matrix, X, which contains the number of times each word was spoken during each time slice (here of length 2.0045 seconds, the TR of the fMRI scan).

The first linear model was estimated separately for each voxel in the fMRI scan using an identity prior on the model weights:

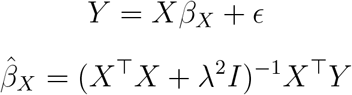

The second linear model was estimated using a semantic prior based on a word embedding space. This embedding space was constructed by computing the statistical co-occurrence of each word in the stories with 985 common English words (see Huth et al., 2016, for details). To apply the semantic prior, the word indicator matrix, *X*, was projected onto the embedding matrix, *E*, and then ridge regression was used to estimate the weights:

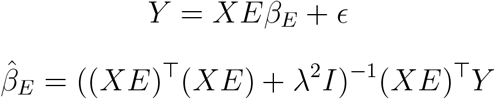

Finally, we used both sets of weights to predict BOLD responses on a separate 10-minute story that had not been used for model estimation, and then computed the correlation between predicted and actual BOLD responses. This model evaluation procedure resulted in two correlation coefficients for each voxel: one for the identity prior and one for the semantic prior. To compare these values we aggregated the data from both subjects, and then computed a 2D histogram of the correlation values (Figure 2).

**Figure 2.**
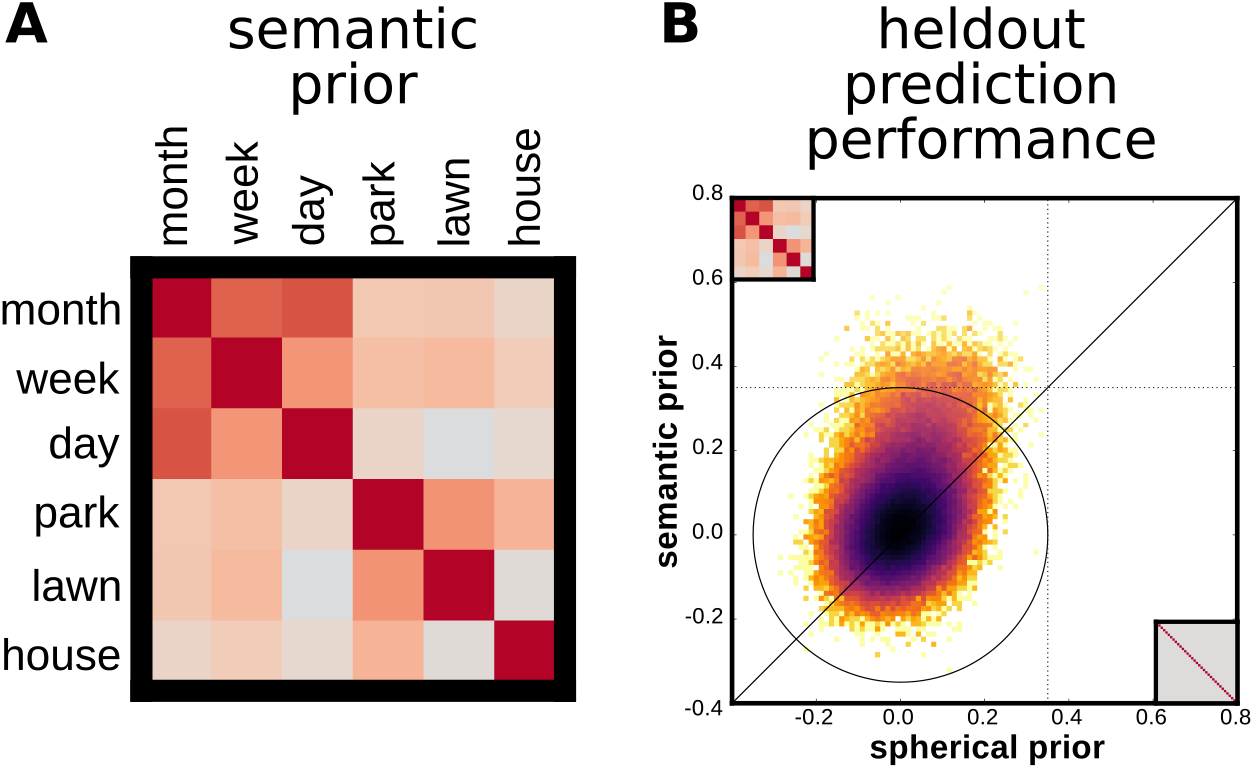
An encoding model estimated using Tikhonov regression and a semantic prior yields more accurate predictions than ridge regression. Two subjects listened to spoken stories while their brain activity was measured with fMRI. Brain activity was modeled as a linear combination of the spoken words in the stories. **(A)** A semantic prior was constructed from word co-occurrence statistics estimated from a large corpus of narrative text. The semantic prior reflects the assumption that words that occur close together are likely to be semantically related. A voxelwise encoding model was estimated using Tikhonov regression and the semantic prior. A separate model was also estimated with a spherical prior (ridge regression). **(B)** 2D histogram across all voxels comparing model prediction accuracy under the semantic prior and the spherical prior. Model prediction accuracy was assessed by computing the correlation between predicted and observed voxel responses to a novel, held-out stimulus. The prediction accuracy was significantly higher (Wilcoxons *W* = 10^9^, *p <* 10^−12^) under the semantic prior (mean pearson *r* = 0.037) than the spherical prior (i.e. ridge regression; mean pearson *r* = 0.005). This suggests that brain responses to the stories are more accurately predicted by encoding model that includes information about the meaning of words (semantics).

Figure 2 shows that model prediction performance is nearly always higher with the semantic prior than with the identity prior, often substantially so. Of approximately 150,000 voxels included in the analysis, about 300 were significantly predicted by the identity prior model, and about 15,000 were significantly predicted by the semantic prior model (n = 291, q(FDR) < 0.05). The difference in model prediction performance is large and significant (Wilcoxon *W* = 10^9^, p < 10^−12^). At worst, we see that some voxels are predicted about as well by both models.

These results suggest that the semantic prior is a much better reflection of the true underlying voxel weights than the identity prior, and thus supports the earlier conclusion that those voxels represent information about the semantic content of language (Mitchell et al., 2008, Wehbe et al., 2014, Huth et al., 2016).

### 3.2 Temporal priors

The temporal activation pattern of the BOLD response is referred to as the hemodynamic response function (HRF). While the neurovascular mechanisms underlying the HRF are not well understood, the shape of the HRF has been extensively studied in humans (Boynton et al., 1996, Glover, 1999). At a first approximation, the neural activation evoked by a stimulus leads to changes in blood-oxygenation that peak 4 to 6 seconds after stimulus onset. Several studies have shown that the shape of the HRF is highly variable across voxels and brain regions both within and across subjects (Aguirre et al., 1998, Handwerker et al., 2004, Kay et al., 2008a). It is important to take this variability into account by estimating the shape of the HRF for each voxel when modeling BOLD responses.

A common approach to estimating the HRF is the use of finite impulse response (FIR) models (Kay et al., 2008a). In an FIR model brain responses are modeled as a linear combination of (*p*) features over a fixed time window (*d*) prior to the stimulus onset (see Equation 1). The number of parameters in an FIR model is much larger (*p × d*) than the original number of features (*p*), and grows linearly with the length of the time window *d*. This increase in the number of parameters can lead to overfitting. In order to reduce overfitting, it is important to regularize FIR models.

When ridge regression is used to estimate FIR models, the implicit assumption is that feature weights are independent across time. This happens because ridge imposes a spherical prior on the temporal covariance of each feature, 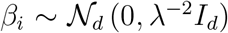 where 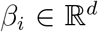 is the vector of weights for feature *i* across the time window. In the Tikhonov framework, we can relax this assumption by specifying temporal priors that are not spherical,

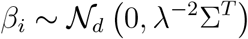

An insight worth highlighting is that applying a temporal prior is equivalent to convolution followed by ridge regression (see Appendix B). This follows from the fact that FIR models can be understood as convolution. In the context of Tikhonov regression, this means that applying a temporal prior of the form *Σ^T^* = (*C^⊤^ C*)^−1^ is equivalent to convolving each feature timecourse with a set of temporal filters given by the columns of *C*^−1^. When *C*^−1^ = *I* the features are convolved with Kronecker delta functions at different delays, which is identical to using delays.

#### 3.2.1 Smoothness temporal prior

One simple and widely studied temporal prior holds that feature weights are smooth across time. This type of prior is typically applied by defining the penalty matrix 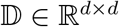 to be a discrete difference operator that penalizes differences between neighboring weights in time,

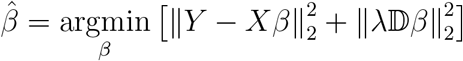

In the Tikhonov framework, this corresponds to a multivariate normal prior with covariance 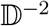 (Wu et al., 2006):

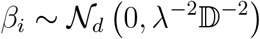

This and similar approaches have been used in several studies (Goutte et al., 2000, Marrelec et al., 2003, Casanova et al., 2008, Bazargani and Nosratinia, 2014).

#### 3.2.2 HRF temporal prior

A more empirically-grounded possibility is to use published mathematical descriptions of the HRF to form a prior (Boynton et al., 1996, Friston et al., 1998, Glover, 1999). Previous work has resulted in the characterization of the commonly used “canonical” HRF. This canonical HRF (*h*_1_), its temporal derivative (*h*_2_), and its derivative with respect to time-to-peak (dispersion; *h*_3_) together provide an informed basis set that can capture some of the empirical variation observed in HRF shapes (Friston et al., 1998). The basis set is a matrix 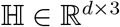

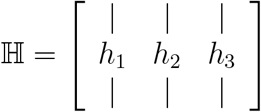

where each *h_i_* is a basis vector of length *d*. However, this basis set is not always flexible enough to capture all voxel- or region-specific variability of the HRF (Woolrich et al., 2004, Kay et al., 2008a, Pedregosa et al., 2015). In such cases, an FIR model with enough statistical power can better estimate the shape of the HRF. In practice, however, the FIR model might be difficult to estimate correctly because the large number of parameters (*p × d*) can lead to overfitting.

Instead of choosing between FIR and HRF-based models, the Tikhohnov framework offers an intermediate approach by allowing us to trade off between both options. To achieve this, we compute the dot product of the HRF temporal basis set and use it as a non-spherical temporal prior on the feature weights,

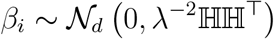

As λ decreases, the effect of the prior on the FIR weights is minimal. On the other hand, as λ increases, the prior has more effect on the FIR weights.

#### 3.2.3 Evaluating temporal MVN priors

In order to evaluate and compare these temporal priors we estimated three encoding models. The first encoding model was estimated with ridge regression, which imposes a spherical temporal prior (Σ^*T*^ = *I_d_*). In the second model, we used Tikhonov regression to impose a smoothness temporal prior on the FIR delays 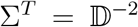. Finally, in a third model we imposed a temporal prior constructed from the covariance of an HRF basis set 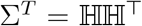 (Friston et al., 1998). All models had the same number of parameters and only differed in the temporal prior used.

We used data from an fMRI experiment in which three subjects watched natural movies while their brain activity was recorded (Huth et al., 2012). A total of 6,555 motion-energy features were extracted from these movies using a three-dimensional Gabor pyramid (Adelson and Bergen, 1985, Watson and Ahumada, 1985, Nishimoto et al., 2011). We used 10 temporal delays in order to account for the HRF (0-20 seconds). This resulted in an FIR model with a total of 65,550 channels and 3,600 time points. We selected the regularization parameter, λ, using a cross-validation procedure (5-fold cross-validation repeated 20 times). This was done separately per voxel for each of the three encoding models estimated. We evaluated model performance for each model by computing the correlation coefficient between predicted and actual BOLD responses on a held-out dataset, which was not used for estimation. The held-out dataset consisted of 270 samples and was constructed by taking the mean temporal BOLD signal across 10 repetitions of a 540 second movie (Schoppe et al., 2016).

A total of approximately 230,000 voxels from three subjects were used in the analyses (Figure 3). We find that the HRF basis set temporal covariance prior provided better predictions than either the spherical prior or the smoothness prior for the best voxels in the population (top 10,000 voxels, Wilcoxon *W* = 10^7^,^24^, *p* < 10^−12^ and *W* = 10^5^,^23^, *p <* 10^−12^, respectively).

**Figure 3.**
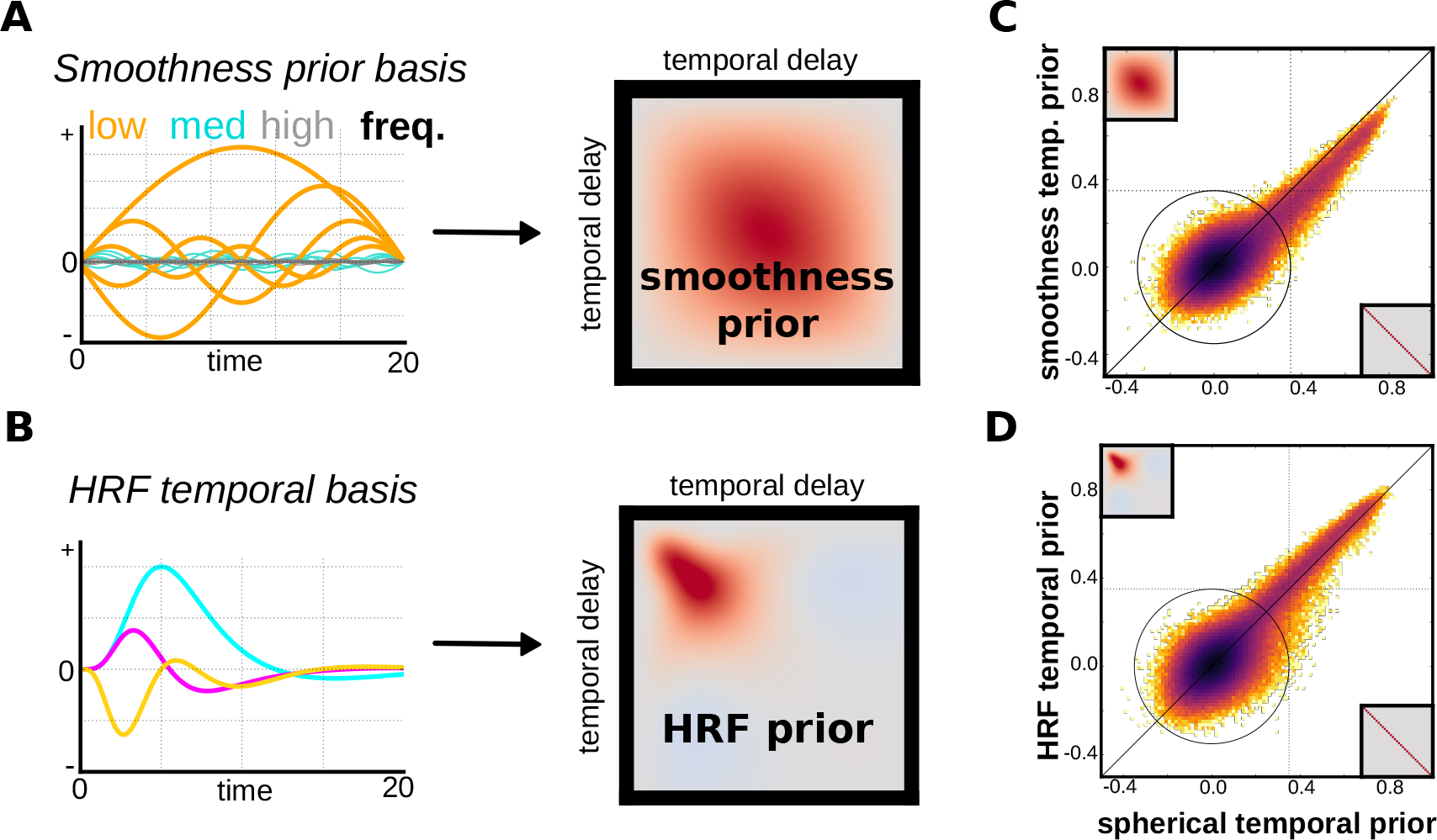
An encoding model estimated using Tikhonov regression and an HRF temporal prior yields more accurate predictions than ridge regression, a smoothness prior does not. Three subjects watched hours of natural movies while their brain activity was measured with fMRI. Three encoding models were estimated each one with a different MVN prior on the covariance of the temporal delays. Two non-spherical temporal MVN priors were compared against a spherical prior (ridge regression). **(A)** The smoothness prior corresponds to a second order difference operator penalty on the temporal delays. This captures the idea that BOLD responses are smoothly varying in time. **(B)** A hemodynamic response function (HRF) temporal MVN prior was built from the temporal covariance of three basis functions. **(C)** The smoothness prior does not improve prediction accuracy on held-out data relative to ridge regression. This is because it enforces high covariance in the middle of the HRF time-course which is an incorrect assumption. **(D)** The HRF temporal basis improves prediction accuracy in well-predicted voxels relative to ridge regression. The improvement in performance is nevertheless small.

The differences in mean prediction performance in the top 10,000 voxels for the models estimated with the HRF (*r* = 0.55 ± 0.001), spherical (*r* = 0.54 ± 0.001) and smoothness (*r* = 0.45 ± 0.001) priors were small but consistent. However, across the total population of voxels the spherical prior yielded better prediction performance (Wilcoxon *W′s* > 10^10^, *p′s* < 10^−12^). These results suggest that for well-predicted voxels at least the HRF prior has a small but consistent advantage.

## 4 Combining spatial and temporal priors

When both a feature prior and a temporal prior are available, they can be used to construct a single spatiotemporal multivariate normal prior. Spatiotemporal priors allow us to incorporate prior information about the covariance of the feature weights and the covariance of the temporal delays when estimating predictive encoding models. However, as the number of features (*p*) and temporal delays (*d*) increase, the spatiotemporal prior matrix becomes large ((*p* × *d*)^2^). This makes the estimation of (non-spherical) spatiotemporal encoding models impractical for neuroimaging. In this section, we present a solution to that makes the estimation of these models tractable when *n < p*.

The spatiotemporal prior is constructed by computing the Kronecker product (⊗) between the feature prior 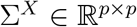 and the temporal prior 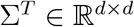,

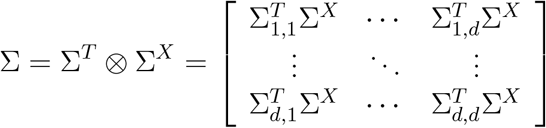

The resulting spatiotemporal prior is 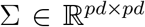. Notice that when both the feature and the temporal priors are spherical, the spatiotemporal prior is also spherical.

The Tikhonov solution to an encoding model with a spatiotemporal multivariate normal prior Σ*^T^* ⊗ Σ^*X*^ can be expressed as (see Appendix C):

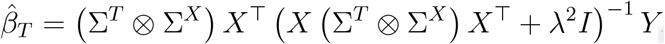

This is equivalent to the ridge regression solution when both priors are spherical (*I_d_* ⊗ *I_p_* = *I_pd_*). However, computing this solution involves constructing an extremely large (*p×d*)^2^ spatiotemporal prior matrix which we would like to avoid. Luckily, the properties of the Kroenecker product allow us to derive a computationally efficient solution in cases where *n < p* (Appendix D). This formulation makes it tractable to fit large encoding models with non-spherical spatiotemporal priors.

### 4.1 Evaluating spatiotemporal MVN priors

To illustrate the power of spatiotemporal priors, we estimated four different encoding models using the data from Huth et al. 2016. We modeled voxel responses to the stimulus as a linear combination of words, and estimated models that differed only in the spatiotemporal prior used:

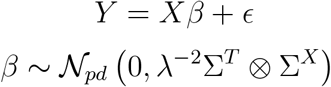

The first and simplest model we evaluated was ridge regression. Ridge regression corresponds to a spatiotemporal prior where both feature and temporal priors are spherical (*I_d_* ⊗ *I_p_*). The second model used a word embedding prior Σ^X^ constructed from word co-occurrence statistics estimated from a large text corpus (described above), and a spherical temporal prior (*I_d_* ⊗ Σ*^X^*). The third model was constructed using a spherical feature prior and a HRF temporal prior Σ*^T^* constructed from a set of HRF basis functions (Σ*^T^* ⊗ *I_p_*). Finally, the fourth model evaluated used a spatiotemporal prior that combines both the word embedding feature prior and the HRF temporal prior (Σ*^T^* ⊗ Σ*^X^*).

The models were constructed using 10 TR temporal delays (20 seconds) in order to account for the hemodynamic lag. A temporal prior 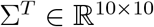 was constructed from the temporal covariance of an HRF basis set during the same time period. The FIR matrix *X* was built using 10 temporal delays for each of the 3,000 channels. This resulted in an FIR feature matrix with a total of 30,000 features and 3,737 time points.

We find that the model estimated with the semantic-temporal prior performs better than the same model estimated with either the semantic or the temporal prior on their own (Figure 4). From a total of about 150,000 voxels, approximately 22,500 were significant (*n* = 291, *q*(*FDR*) < 0.05) when using the semantic-temporal prior (*r* = 0.045 ± 0.0003), approximately 5,500 with the temporal prior (*r* = 0.019 ± 0.0002), and 15,000 with the semantic prior (*r* = 0.037±0.0002). The semantic-temporal prior performs much better than the temporal prior model alone (Wilcoxon *W* = 10^9.67^, *p* < 10^−12^). This is not surprising since the semantic-temporal prior includes the semantic prior and that on its own improves prediction performance (see Figure 2). However, we find that the semantic-temporal prior improves performance over and above the semantic prior alone (Wilcoxon *W* = 10^9.71^, *p* < 10^−12^). In sum, we can gain the best from both worlds by combining feature and temporal priors into a single spatiotemporal prior and thereby improve the prediction performance of encoding models.

**Figure 4.**
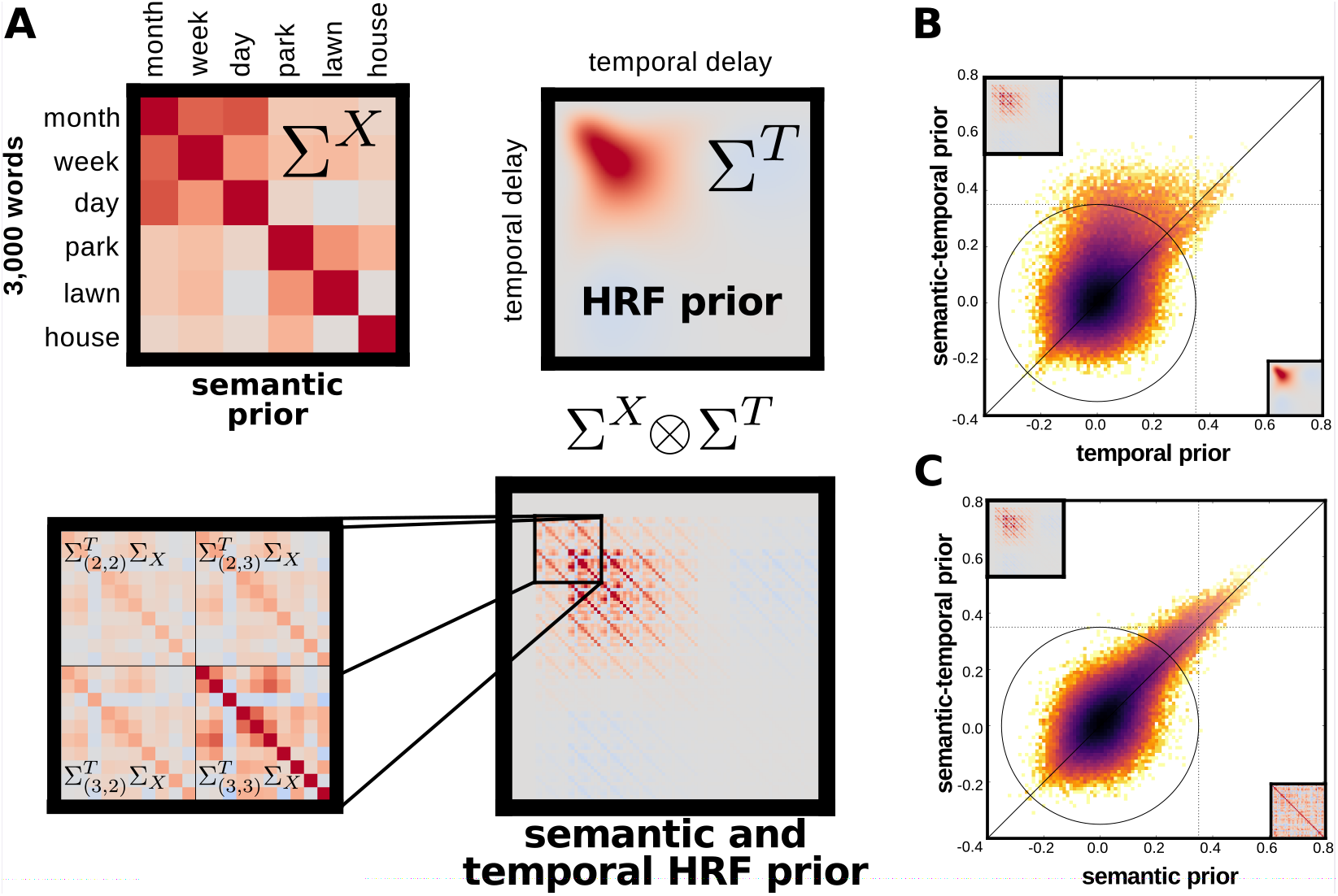
An encoding model estimated using Tikhonov regression and a single spatiotemporal prior yields more accurate predictions than using a temporal or a feature prior alone. Two subjects listened to spoken stories while their brain activity was measured with fMRI. Brain activity was modeled using Tikhonov regression and a spatiotemporal MVN prior built by combining an HRF temporal prior and a semantic prior. **(A)** The spatiotemporal MVN prior is constructed by computing the Kroenecker product of the HRF temporal and semantic priors. This is equivalent to scaling the semantic prior by each element in the HRF temporal prior and concatenating the resulting matrices. **(B,C)** The spatiotemporal MVN prior consistently yields better prediction accuracy than using either the HRF temporal or the semantic priors on their own.

## 5 Combining spatiotemporal priors

It is becoming increasingly important to characterize how and where different feature spaces overlap in terms of variance explained (Borcard et al., 1992, Lescroart et al., 2015, de Heer et al., 2017). This is usually done by combining different feature spaces into one single joint model (Lescroart et al., 2015, Çukur et al., 2016, de Heer et al., 2017). However, estimating joint models with ordinary regularization techniques (e.g. ridge, LASSO, elastic net) assumes that all feature spaces require the same level of regularization. This is often an incorrect assumption. In practice, the level of regularization for a feature space depends on factors such as the feature space covariance, the number of features, and the fraction of variance explained by that feature space. The choice of regularization level for each feature space is critically important to the prediction accuracy of models that combine multiple feature spaces.

Suppose we have two feature spaces 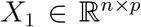 and 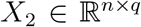 that are combined into a single encoding model:

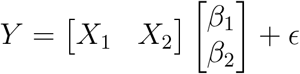

Using ridge regression to estimate such a model is equivalent to choosing the same level of regularization on each feature space. The ridge prior on the joint feature weights can be expressed as:

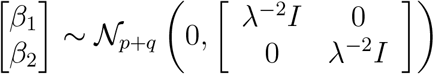

where the regularization level λ is selected via cross-validation or other methods. It is clear that the prior on each feature space is the same (λ^−2^*I*). However, feature spaces *X*_1_ and *X*_2_ might need different levels of regularization. Estimating joint models with the same level of regularization on each feature space can lead to poor prediction performance. This is because the globally optimal λ will often be suboptimal for the individual feature spaces. This issue applies to ordinary ridge regression (Hoerl and Kennard, 1970), LASSO (Tibshirani, 1996), and elastic-net (Zou and Hastie, 2005) models.

### 5.1 Banded ridge regression

Instead, we can impose separate priors on the weights for each feature space:

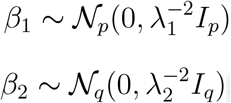

The Tikhonov framework allows us estimate the joint model with a separate prior on each feature space,

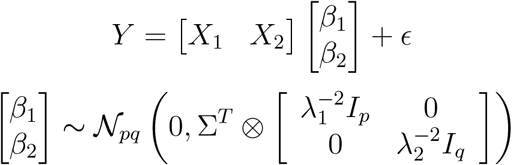

where λ_1_ and λ_2_ can take different values. For the sake of clarity, assume a spherical temporal prior (Σ^T^ = *I_d_*). Estimating this model is equivalent to solving:

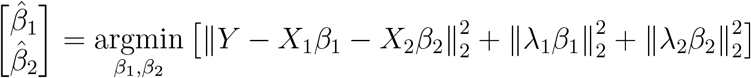

The solution is:

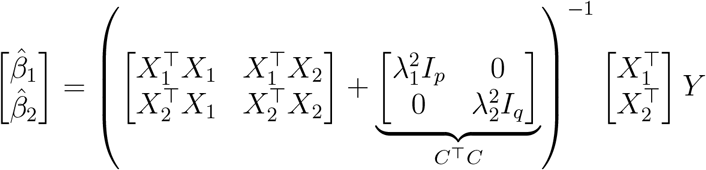

Notice that the penalty (*C*^T^*C*) becomes the ridge penalty when when λ_1_ = λ_2_. However, when λ1 ≠ λ2 the structure of the penalty becomes “banded” with the first *p* values along the diagonal equal to λ_1_ and the next *q* values equal to λ_2_.

We can also transform the Tikhonov problem into standard form:

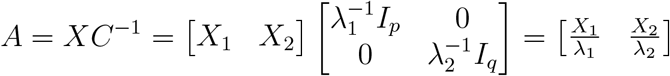

This is a surprisingly simple expression. It says that scaling the features is equivalent to adjusting the strength of the prior. This is due to the inverse relationship between feature scaling and feature weights. All else being equal, dividing the features by a constant is equivalent to multiplying the weights by that constant. Finally, the kernelized standard form solution becomes

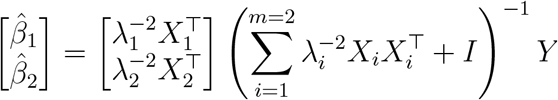

### 5.2 Evaluating banded ridge regression

We evaluated banded ridge regression using data from a natural movie experiment (Huth et al., 2012). We constructed a single encoding model that combined two previously published feature spaces. The first feature space 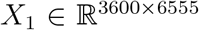 captured low-level visual properties from the stimulus (Nishimoto et al., 2011). The second feature space 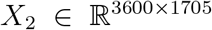 captured high-level visual properties consisting of object and action categories (Huth et al., 2012). We modeled voxel responses as a linear combination of these feature spaces:

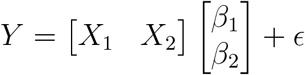

We estimated this joint model using standard ridge regression and banded ridge regression. The only difference between these models is the feature prior used: ridge regression uses a spherical prior 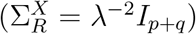 whereas banded ridge regression uses a non-spherical prior:

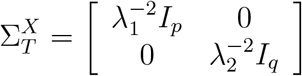

Low-level motion-energy features were extracted from the natural movies using a three-dimensional Gabor pyramid (Adelson and Bergen, 1985, Watson and Ahumada, 1985, Nishimoto et al., 2011). This yielded a total of 6,555 features which differed in orientation, spatial and temporal frequency, location, size, and direction of motion. The high-level object and action category features were tagged by hand from each one second segment of the movies and labeled using WordNet synsets (Miller, 1995, Huth et al., 2012). The hyponyms for each synset were inferred from the WordNet graph and also included. This process yielded a total of 1,705 object and action category features. An FIR model was then constructed by including 10 TR temporal delays for each feature in order to account for the hemodynamic response function. The resulting model consisted of 8,260 stimulus features times 10 delays (82,600 total features) and 3,600 time points. For simplicity, we used a spherical temporal prior. The feature prior hyperparameters for both ridge (λ) and banded ridge (λ_1_ and λ_2_) models were selected per voxel via 5-fold cross-validation. The performance of each model was assessed by computing the correlation between model predictions and actual responses using a held-out dataset not used for model estimation.

Banded ridge regression provided far better joint model predictions than standard ridge regression (Figure 5; Wilcoxon *W* = 10^9 92^, *p* < 10^−12^). Of the approximately 230,000 voxels, about 40,000 were significantly predicted with banded ridge (*n* = 270, *q*(*FDR*) < 0.05, *r* = 0.06 ± 0.0003). In contrast, approximately 20,000 voxels were significantly predicted with ridge regression (*n* = 270, *q*(*FDR*) < 0.05, *r* = 0.3 ± 0.0002). We used the estimates from the banded ridge joint regression to compute the prediction performance of each feature space on its own. This gives us a separate prediction performance value per voxel for each the motion-energy features and for the object category features. These prediction performance values are plotted on the cortical sheet in Figure 5C. There is a strong separation in prediction performance between early visual cortex being best predicted by motion-energy features, and higher visual cortex better predicted by object category features.

**Figure 5.**
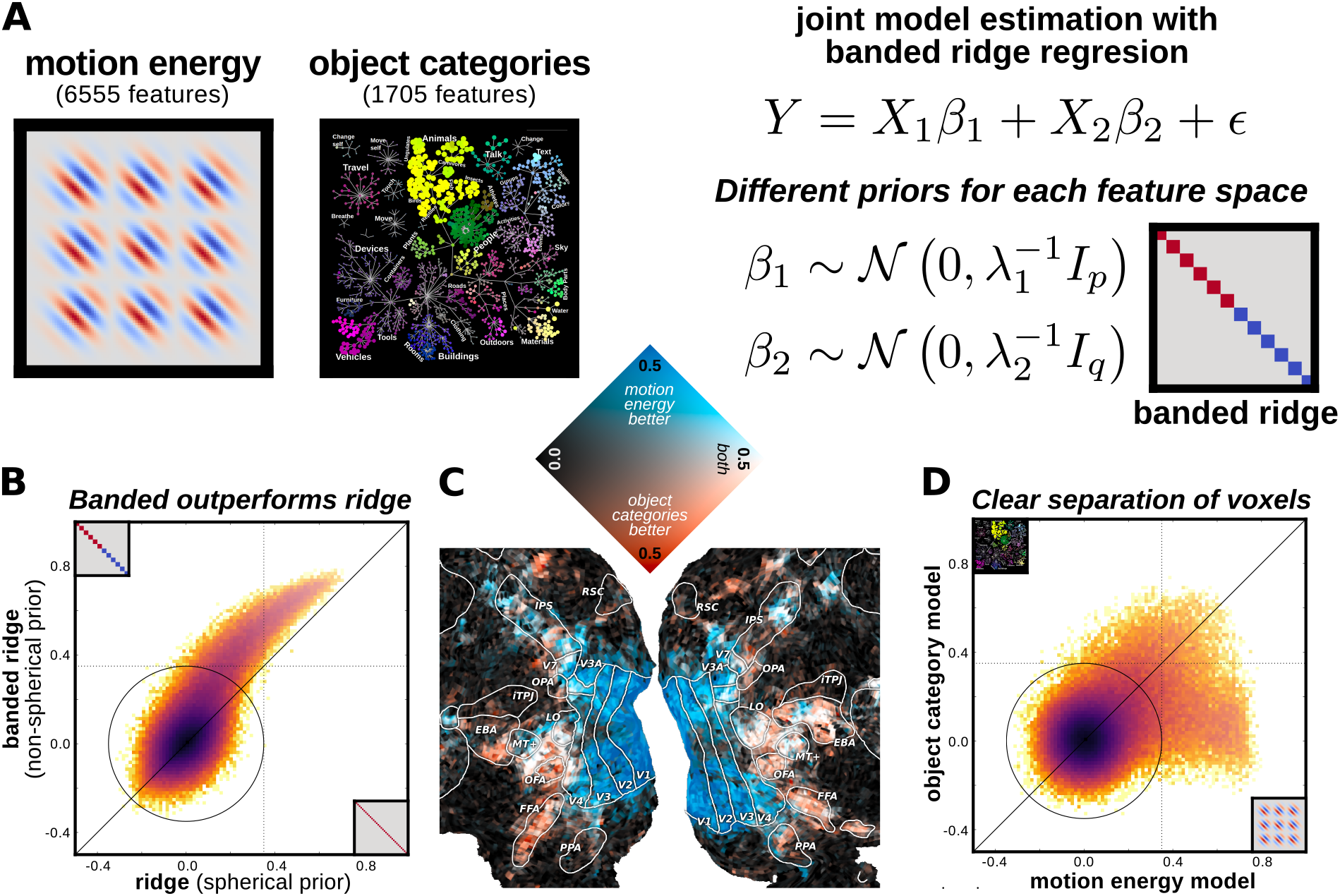
Banded ridge regression yields more accurate predictions than ridge regression when estimating joint models. Three subjects watched natural movies while their brain activity was measured with fMRI. **(A)** Brain activity was modeled as a linear combination of motion energy and object category features. A joint model was estimated using Tikhonov regression and a non-spherical MVN prior built by combining two spherical MVN priors, one for each feature space. We refer to this approach as banded ridge regression. **(B)** Banded ridge regression yields higher prediction accuracy on held-out data than ridge regression (Wilcoxons W = 10^9.92^, p < 10^−12^). **(C)** Once the joint model is estimated with banded ridge, the prediction accuracy of the individual feature spaces can be computed. Prediction accuracy for each feature space is plotted simultaneously on the cortical sheet of one subject with a 2D colormap. Voxels in early visual cortex are well-predicted by motion-energy features (blue) and voxels in higher visual cortex are well-predicted by object category features (red). A subset of voxels are predicted similarly well by both models (white). **(D)** 2D histogram shows the separation of voxels well fit by each of the two models. A large set of voxels are better predicted by motion-energy features than by object category features. Conversely, many voxels are better predicted by object category features than by motion-energy features.

A big benefit of banded ridge regression is that it removes spurious correlations between feature spaces. When the motion-energy model is estimated by itself with ridge regression, even voxels in higher visual cortical regions can be well-predicted (Figure 6A). This can occur because of stimulus correlations. For example, suppose there is a consistent correlation between between vehicles and left-and right-direction selective motion-energy filters in the lower visual field. Estimating the motion-energy model on its own will yield high predictions. By estimating the object category and motion-energy models together, the variance can be correctly assigned to the object category model. In cases where a close to perfect correlation exists, the banded ridge estimation will split the variance among the feature spaces. In sum, banded ridge regression yields better estimates of the variance that can be explained by any one feature space.

**Figure 6.**
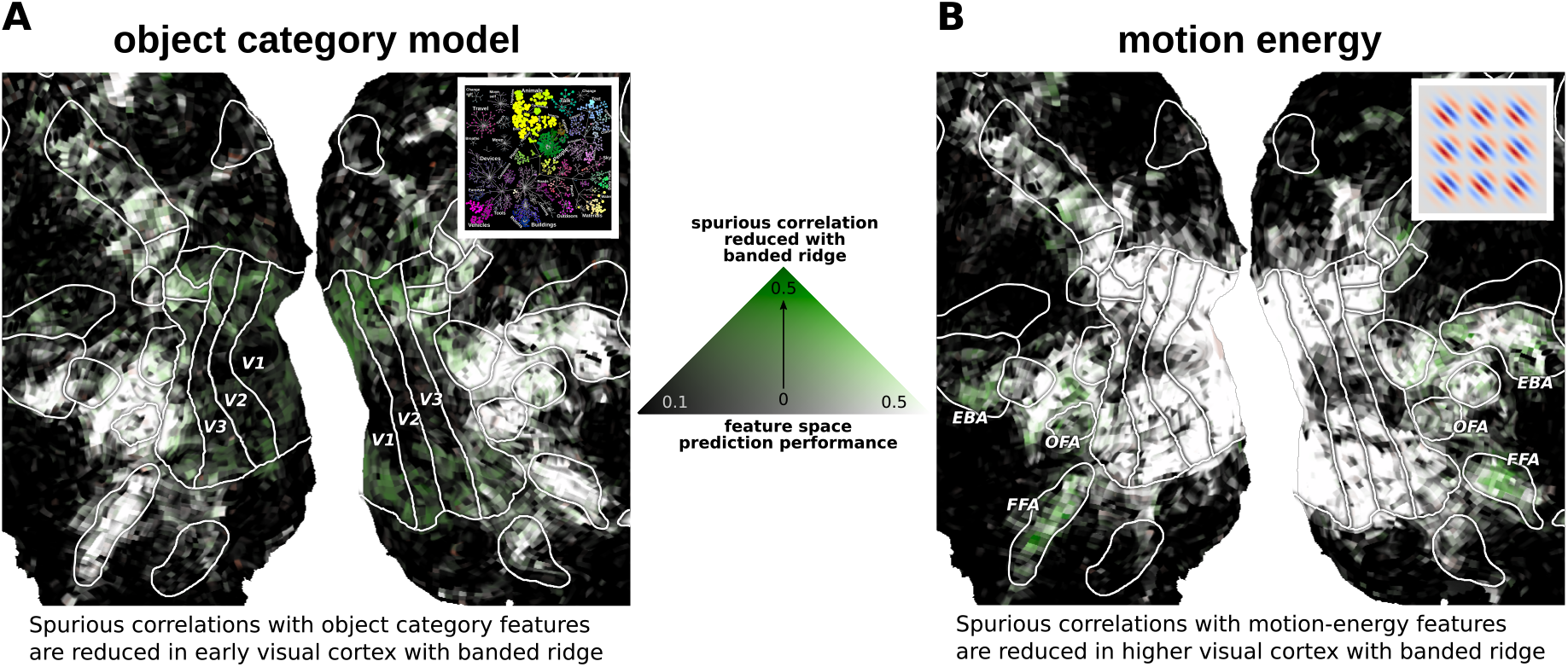
Banded ridge regression reduces spurious prediction accuracy. (A) Ridge regression was used to model brain activity as a linear combination of object category features. Separately, banded ridge regression was used to model brain activity as a linear combination of both object category and motion energy features (see Figure 5). The prediction accuracy of object category features was computed for both the banded ridge and the ridge regression models separately. The prediction accuracy of each model is plotted simultaneously on the cortical sheet of one subject with a 2D colormap. Voxels located in anterior visual cortex are accurately predicted by object category features under both standard ridge regression and banded ridge (voxels colored in white). However, voxels located in early visual cortex (EVC) appear to be better predicted by object category features under standard ridge regression than under banded ridge. This difference occurs because ridge regression attributes all variance to object category features, while under banded ridge regression the variance is split correctly between object category and motion energy features. Because banded ridge regression allows motion energy features to explain some of the variance away from object category features, it reduces the artifactual predictive power of object category features observed under ridge regression in EVC voxels (colored in green). **(B)** The prediction accuracy of motion energy features was computed for both the banded ridge and the ridge regression models separately. Voxels located in EVC are accurately predicted by motion energy features with both standard ridge regression and banded ridge (voxels colored in white). Banded ridge regression allows object category features to explain some of the variance away from the motion energy features in anterior visual cortex voxels (colored in green).

## 6 Discussion

The results highlighted in this paper show that the Tikhonov framework works very well for estimating predictive models of BOLD responses in the context of naturalistic experiments. The Tikhonov framework can be used to incorporate prior information about how features in the model covary, how the measured signals vary in time, and how multiple feature spaces can be used to build a single predictive model.

The reader should be aware that our results do not necessarily generalize to every experimental condition or dataset. In general, the experimenter should treat the choice of fitting procedure (e.g. FIR, grouped L1, OLS, etc) as a hyperparameter on its own right and use statistical learning theory to make a decision. The Tikhonov framework is presented as another method in the toolkit available to researchers. The banded ridge model proposed is of particular utility when estimating joint models that combine several feature spaces to predict brain activity. We have shown a computationally efficient framework for incorporating multivariate normal priors into spatiotemporal encoding models. And that this framework is flexible enough to work well in a variety of cases. The software used to estimate all the models presented in this paper is publicly available (http://github.com/gallantlab/tikreg). We hope this facilitates the adoption of this framework.

## Acknowledgments

This work was supported by grants from the National Science Foundation (NSF; IIS1208203), the National Eye Institute (EY019684 and EY022454) and the Office of Naval Research (N00014-15-1-2861). A.G.H. was also supported by a Burroughs-Wellcome fellowship. We thank Leila Wehbe for useful technical discussions, and Brittany Griffin for segmenting and flattening cortical surfaces. The authors declare no conflict of interest.

## Appendix A Standard form derivation

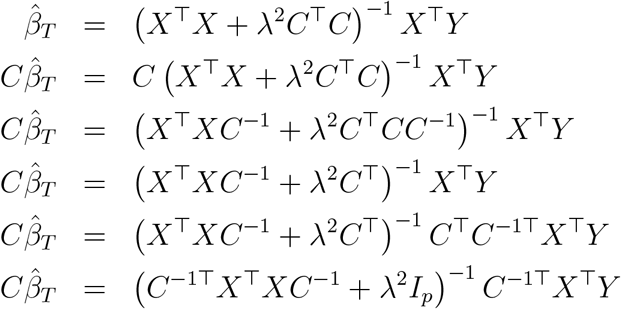

Define *A = XC*^−1^ 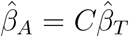. The solution becomes

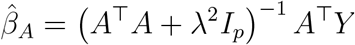

and one can recover the original weights with

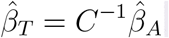

There exists an interesting relationship between the prior covariance matrix, Σ, and the Tikhonov penalty matrix, *C*. When the penalty Gram matrix, *C*^T^*C*, is full-rank, it is invertible and there exists a corresponding prior,

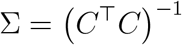

However, the standard form decouples the two concepts. There exist well-defined Tikhonov penalties for which a prior cannot be expressed. In particular, if the penalty Gram matrix is not positive semi-definite, no inverse exists and therefore the prior cannot be formally expressed. The converse is also true. A rank-deficient prior can be used if the problem is in standard form, yet there is no corresponding penalty matrix. See Doicu et al. (2010) for a full treatment.

## Appendix B Equivalence of FIR models with temporal priors and convolution followed by ridge

Estimating an FIR model with a temporal prior 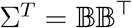

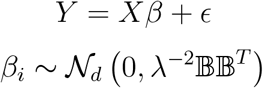

is equivalent to convolving the features xi with the columns of 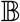 and estimating the model using ridge regression:

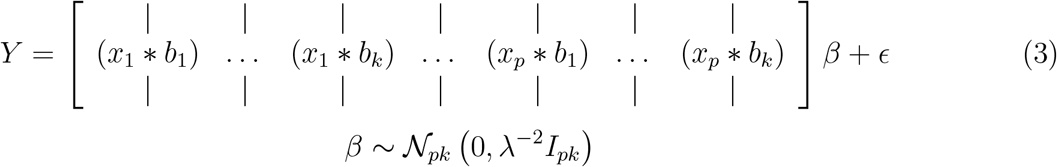

Recall the definition of the standard form transform:

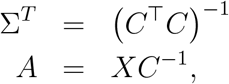

where 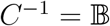 for a temporal prior 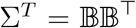. The standard transform of the FIR model can be written as

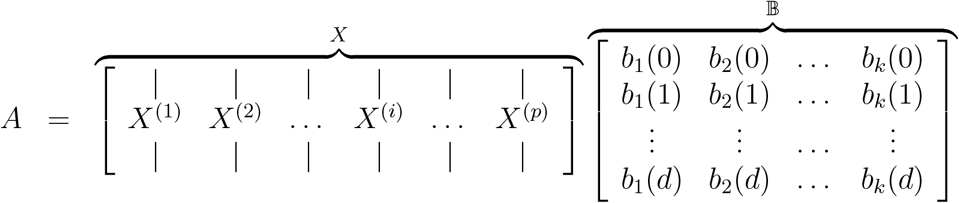

where each 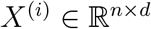 is a matrix that contains every feature 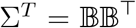 at delays 0 through *d*, and every row of *X*^(*i*)^corresponds to a particular time point *t*:

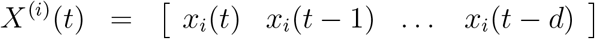

We can express every entry of the matrix A as the dot product between *X*^(*i*)^(*t*) and each column of the temporal basis set, *bj*:

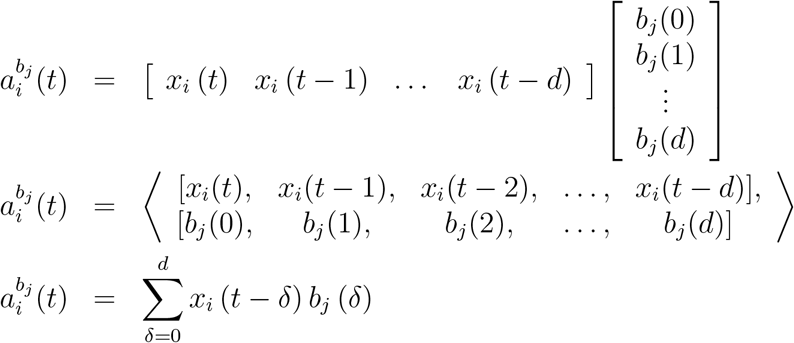

which is the definition of discrete convolution

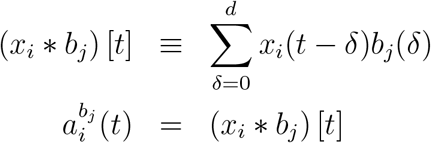

Finally, we rewrite 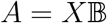 as the convolution of each feature *i* with each temporal basis *j*

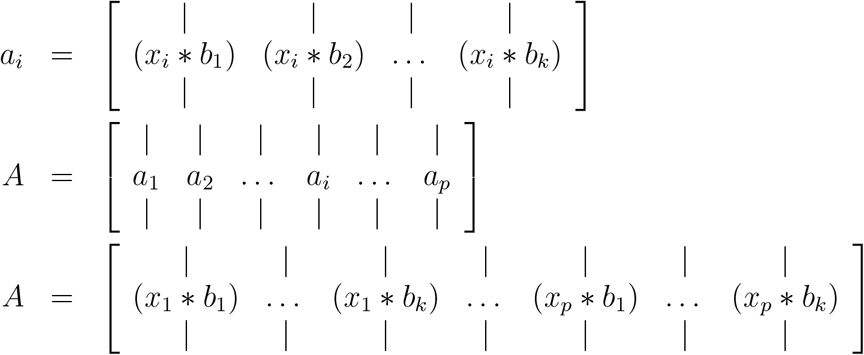

This is exactly Equation 3.

## Appendix C Kernel solution to encoding models with spatiotemporal MVN priors

The standard form solution is

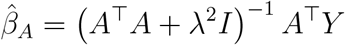

The kernel solution to the standard form problem becomes

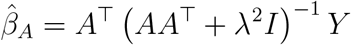

Expanding this out using the fact that *A* = *XC*^−1^

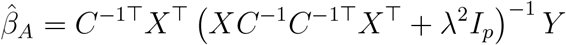

We know Σ = C ^1^C ^1T^ = (CC^T^) ^1^. Replacing this in

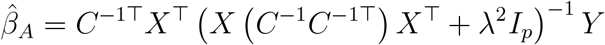

To recover the Tikhonov solution, recall that β_T_ = C ^1^,β_A_. Substituting this in

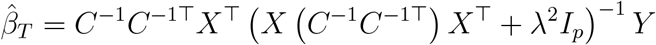

We know Σ = *C*^1^*C*^1T^. Replacing this in

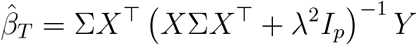

In the case of spatiotemporal kernels Σ = Σ^*T*^ ⊗ Σ^*X*^. The full solution becomes

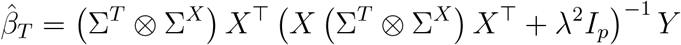

## Appendix D Efficient kernel solution for models with spatiotemporal MVN priors

We now derive a computationally efficient solution for the kernel solution for an encoding model with non-spherical spatiotemporal multivariate normal priors. This formulation makes the estimation of these models computationally tractable.

The spatiotemporal prior is constructed by computing the Kronecker product (⊗) between the feature prior 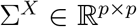 and the temporal prior 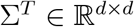,

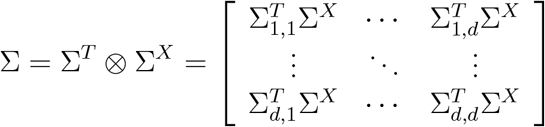

The resulting spatiotemporal prior is 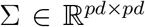. Notice that when both the feature and the temporal priors are spherical, the spatiotemporal prior is also spherical.

The Tikhonov solution to an encoding model with a spatiotemporal multivariate normal prior Σ^*T*^ ⊗ Σ^*X*^ can be expressed as (see Appendix C):

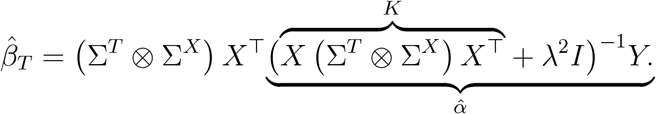

A computationally efficient solution can be derived by re-arranging terms. First, notice that the kernel regression solution to the standard form problem is embedded within the Tikhonov solution above:

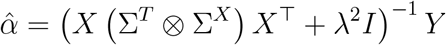

The term inside the parenthesis is the regularized *n* × *n* kernel matrix K of the standard form transformation:

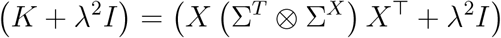

Recall that *X* is an *n* × *pd* FIR matrix which includes delayed copies of the linearized stimulus feature matrix. Computing the kernel matrix thus requires the following matrix

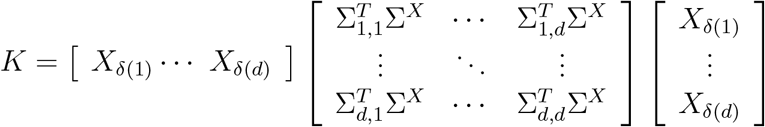

Finally, this matrix multiplication can be expressed as a sum of matrix products,

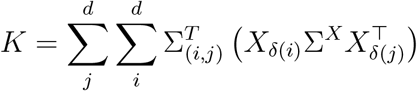

This formulation makes the problem of estimating encoding models with spatiotemporal multivariate normal priors tractable in contexts when *n* < *p*.

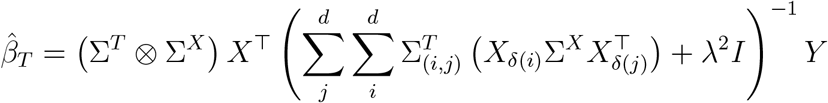

## Appendix E

Extension to priors on priors: hyper-priors

We have shown the usefulness of imposing various temporal and spatial priors on feature weights to improve predictive models. There exist situations, however, when the expert prior itself needs to be regularized. This can be the case when the expert prior is derived empirically and is noisy, or when the prior can be modified to match the data better. In such cases, we can apply the same principle and impose a prior on the prior—a hyper-prior. We next show an example on how to incorporate hyper-priors to our framework.

In the section on Temporal Priors we described the smoothness prior. Our results show that imposing a smoothness prior on the temporal delays does not improve the prediction performance of the motion-energy model. This is surprising. We expect the haemodynamic response function to be temporally smooth, and so imposing a smoothness prior should improve prediction performance. This intuition, however, ignores the structure of the smoothness prior.

The smoothness prior imposes a strong covariance to delays in the middle of the temporal filter (see Figure 3). This is problematic because the goodness of the prior will depend on the number of delays. This is a bad assumption in many cases. In order to avoid this issue, we can impose a spherical prior on the smoothness prior. This can be thought of as trading off between a spherical prior and the smoothness prior, where the tradeoff is controlled by the hyper-prior hyper-parameter.

In general, hyper-priors can be expressed as

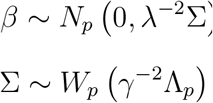

where *W* is a Wishart distribution. In the case of the smoothness prior, this results in

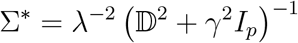

where λ and γ are hyper-parameters.

Estimating models that include both a prior and a hyper-prior is feasible under our framework (and implemented in the accompanying software). However, this flexibility comes at the cost of computational resources because the hyper-prior hyper-parameter (γ) needs to be estimated via cross-validation.

